# A warmer growing season triggers earlier following spring phenology

**DOI:** 10.1101/2021.08.08.455549

**Authors:** Hongshuang Gu, Yuxin Qiao, Zhenxiang Xi, Sergio Rossi, Nicholas G. Smith, Jianquan Liu, Lei Chen

## Abstract

Under global warming, advances in spring phenology due to the rising temperature have been widely reported. However, the physiological mechanisms underlying the warming-induced earlier spring phenology remain poorly understood. Here, using multiple long-term and large-scale phenological datasets between 1951 and 2018, we show that warmer temperatures during the previous growing season between May and September led to earlier spring phenology in the Northern Hemisphere. We also found that warming-induced increases in maximum photosynthetic rate in the previous year advanced spring phenology, with an average of 2.50 days °C^-1^. Furthermore, we found a significant decline in the advancing effect of warming during the previous growing season on spring phenology from cold to warm periods over the past decades. Our results suggest that the observed warming-induced earlier spring phenology may be driven by increased photosynthetic carbon assimilation in the previous season, while the slowdown in the advanced spring phenology arise likely from decreased carbon assimilation when warming exceeding the optimal temperatures for photosynthesis. Our study highlights the vital role of photosynthetic carbon assimilation during growing season in spring phenology under global warming.

## Introduction

Plant phenology influences the fitness of individual plants and functioning of terrestrial ecosystems, including the fluxes of water and energy and food webs^1–6^. Since phenological events are highly sensitive to climate variations, monitoring changes in plant phenology can provide the first clear visible signals of the impact of climate change on terrestrial ecosystems^6,7^. Under global warming, advanced spring phenology due to rising temperature has been widely reported^8–12^. However, important questions regarding the physiological mechanisms underlying this response remain unanswered^13–17^. This largely hinders the prediction of spring phenology and global carbon cycling under future warming conditions.

Generally, spring phenology is considered to be driven by temperatures in winter and spring because plants need to accumulate sufficient winter chilling to end endodormancy and spring forcing units to break ecodormancy before spring phenology ^18–22^. Recent studies show that the response of earlier spring phenology to climate warming is declining^17^. However, there continues to be debate about the drivers of the slowdown in the warming-induced spring phenology. In fact, plants need to assimilate and store sufficient carbohydrates in the preceding growing season to resist to the frost temperatures in winter and support growth reactivation in spring^23–26^. In temperate regions, nonstructural carbohydrates (NSC; soluble sugar and starch) often reach the maximum levels in autumn before winter dormancy, but become depleted by early summer after spring growth^27–29^. Girdling experiments have demonstrated that a later budbreak is often associated with a lower NSC availability^30,31^. The timing of spring phenology is therefore likely to depend on the photosynthetic carbon assimilation during the previous growing season.

Under global warming, increasing temperatures may influence the photosynthetic carbon assimilation and alter spring phenology in the following year ^32^. Photosynthetic carbon uptake tends to show a peaked response to temperature at leaf and canopy scale^12,33–36^. As such, an increase in temperature might increase photosynthesis in cold and temperate regions, and advance spring phenology^37,38^. When temperatures increase above the optimal threshold for photosynthesis, this could explain the slowdown in warming-induced advancement in spring phenology. However, previous researches have largely overlooked the effect of previous growing season climate on spring phenology^39–42^.

Using long-term phenological observations and remote-sensing chronologies collected in the Northern Hemisphere (Fig. 1), we analyzed the effect of warming during the previous growing season on spring phenology. We hypothesized that timing of spring phenology may depend on the photosynthetic carbon assimilation during the previous growing season prior to leaf senescence. According to this carbon-driven assumption, warmer temperatures during the previous growing season are expected to increase photosynthetic carbon uptake and trigger earlier spring phenology.

**Fig. 1.**
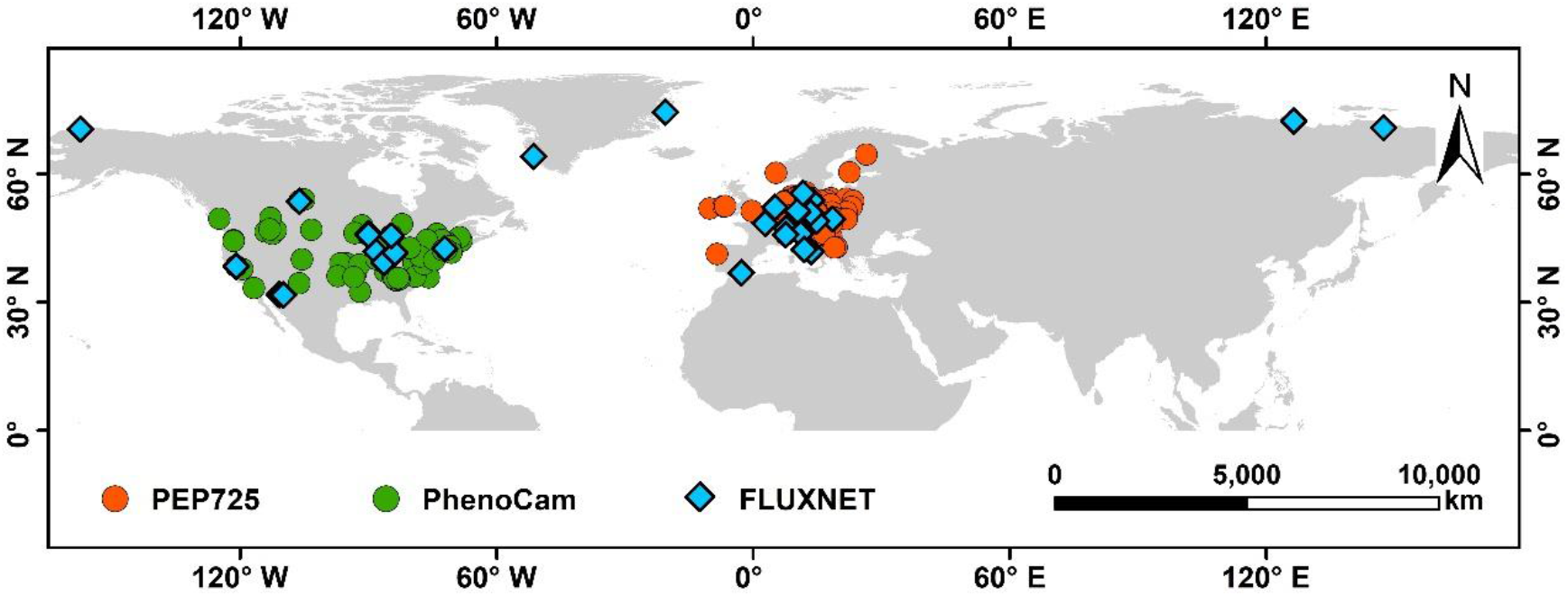
Distributions of the phenological observation sites in this study. Orange dots represent the 2,300 sites selected from the PEP725 dataset across central Europe. Green dots and blue diamonds represent 78 sites in North America from the PhenoCam network and 39 FLUXNET sites, respectively.

## Materials and Methods

### PEP725 phenological network

Data were provided by the European phenology database PEP725 (http://www.pep725.eu/), which contains phenological observations of temperate species across central Europe since 1951^43^. We selected the date when the first leaf stalks were visible (BBCH 11 in PEP725) to represent the start of spring phenology (SOS) and date when 50% leaves had their autumnal color (BBCH94 in PEP725) to represent the end of autumn phenology (EOS). Data exceeding 2.5 times of median absolute deviation (MAD) were considered outliers and removed^44^. We selected 466,988 records of nine temperate tree species (Table S1) at 2,300 sites, for a total of 171,202 species-site combinations with at least 30-year observations.

### PhenoCam network

The PhenoCam network (https://phenocam.sr.unh.edu/) is a cooperative database of digital phenocamera imagery which provides the dates of phenological transition between 2000 and 2018 worldwide^45,46^. In the PhenoCam network, the 50%, and 90% of the Green Chromatic Coordinate (G_CC_) were calculated daily to extract the date of greenness rising and falling based on the following formula:

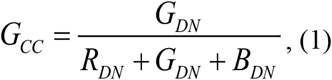

where *R*_*DN*_, *G*_*DN*_ and *B*_*DN*_ are the average red, green and blue digital numbers (DN), respectively.

We selected 50% threshold of G_CC__90 (G_CC_ reaches 90^th^ quantiles of its seasonal amplitude) as SOS^47^. We removed outliers according to the above-mentioned procedure, and we selected sites with at least 8-year observations between 2000 and 2018. We also excluded agricultural ecosystems to avoid human influence. The final dataset had a total of 738 records at 78 sites from three vegetation types: deciduous broadleaf forests, evergreen forests and grassland.

### GIMMS NDVI_3g_ phenological product

The Normalized Difference Vegetation Index (NDVI), a proxy of vegetation greenness and photosynthetic activity, is commonly used to derive phenological metrics^48^. We derived SOS from the third generation GIMMS NDVI_3g_ dataset (http://ecocast.arc.nasa.gov) from Advanced Very High Resolution Radiometer (AVHRR) instruments for the period 1982-2014 with a spatial resolution of 8 km and a temporal resolution of 15 days^49^.

We only kept areas outside tropics (latitudes >30 °N), which have a clear seasonal phenology^50^ and excluded bare lands with annual average NDVI < 0.1 to reduce bias. We applied a Savizky-Golay filter^51^ to smooth the time series and eliminate noise of atmospheric interference and satellite sensor, and used a Double Logistic 1^st^ to extract phenology dates^50^ according to the formula:

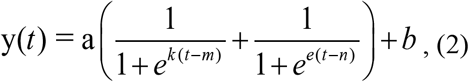

where a, *k, m*, and *n* are parameters of logistic function and a is the initial background NDVI value, a + *b* represents the maximum NDVI value, *t* is time in days, and y(*t*) is the NDVI value at time *t*. The second-order derivative of the function (Eq. (2)) was calculated to extract SOS and EOS at the first and second local maximum point, respectively^52,53^.

### FLUXNET dataset

The flux dataset was downloaded from FLUXNET (https://fluxnet.org/data/). The data were released in November 2016 (total 212 sites) worldwide^54^. The dataset was processed with a processing pipeline to reduce uncertainty by improving the data quality control. The pipeline generates uniform and high-quality derived data products suitable for studies that compare multiple sites^54^. We selected 39 sites with at least 5-year observations and daily records > 300 for each year between 1992 and 2014. The Singular Spectrum Analysis (SSA) filter method^55^ was used to smooth the time series of gross primary productivity (GPP) to minimize the noise. GPP_max_, daily maximum GPP in a year, is considered as an important index to evaluate the carbon fixation of terrestrial ecosystems and the feedback of vegetation climate^56–58^. We extracted the GPP_max_ from the smoothed GPP curve by the SSA-based de-nosing smoothing method^59^. SOS and EOS were extracted from smoothed daily GPP curve based on the threshold method^51^. The spring and autumn threshold were defined as 15% of the multi-year daily GPP maximum following previous studies^60,61^, and SOS and EOS were defined as the turning point when the smoothed GPP was higher or lower than spring or autumn threshold, respectively.

### Climate data

Gridded daily mean temperature, precipitation, solar radiation and air humidity during 1950-2015 in Europe were downloaded from the database E-OBS (http://www.ecad.eu/)^62^ at 0.25° spatial resolution. Gridded monthly soil moistures during 1979-2015 were downloaded from World Meteorological Organization (http://climexp.knmi.nl/select.cgi?id=someone@somewhere&field=clm_wfdei_soil01) at 0.5° spatial resolution and banded with PEP725 dataset. Global monthly mean temperatures during 1981-2017 were downloaded from Climate Research Unit (https://crudata.uea.ac.uk/cru/data/hrg/cru_ts_4.04/cruts.2004151855.v4.04/) at 0.5° spatial resolution and matched the PhenoCam and GIMMS NDVI_3g_ datasets. Bilinear interpolation method was used to extract climate data of each site or pixel using the *raster* package^63^ in R version 4.0.3^64^. Environmental variables, including daily mean temperature (°C), shortwave radiation (Wm^-2^), CO_2_ (ppm), and precipitation (mm) of FLUXNET dataset were also extracted.

### Statistical analysis

To tested our hypotheses, we primarily used the observations from the PEP725 network corresponding E-OBS climate dataset. We did this because PEP725 data was relatively more reliable than the extracted phenological metrics from imagines of PhenoCam network and GIMMS NDVI_3g_ product because its phenological records were taken manually *in situ*. In addition, the PEP725 network covered a longer period (between 1951 and 2015) than PhenoCam (between 2000 and 2018) and GIMMS NDVI_3g_ dataset (between 1982 and 2014). The PhenoCam and GIMMS NDVI_3g_ phenology products were used to test the robustness and generality of the results obtained from the PEP725 network. Specifically, we calculated the temperature sensitivity (S_T_, change in days per degree Celsius) based on mean temperatures during the previous growing season from May to September (T_GS_) and timing of spring phenology using three complementary large-scale datasets (PEP725, PhenoCam, GIMMS NDVI_3g_) in the Northern Hemisphere. To clarify the underlying physiological mechanisms, we further examined the relationships between GPP_max_ of previous growing season and SOS between 1992 and 2014 using FLUXNET data.

### Temperature sensitivities

Temperature sensitivity (S_T_, change in days per degree Celsius), defined as the slope of a linear regression between the dates of phenological stages and the temperature^21,65,66^, was used to investigate the effects of T_GS_ on leaf unfolding dates in the PEP725 network. The length of growing season was defined as the period between SOS and EOS. The mean dates of SOS and EOS from the PEP725 network were DOY 120 and DOY 280. Therefore, the period between May and September was selected to represent the growing season. Linear regression models were used to calculate S_T_ of leaf unfolding for each species at each site. In the model, the response variable was the leaf unfolding date while the predictor was the T_GS_.

In addition, a linear mixed-effects model was used to exclude the co-variate effects of other climate factors and autumn phenology, and further examine the overall effect of T_GS_ on leaf unfolding dates by pooling all records across species and study sites. In the model, the response variable was leaf unfolding dates, and the predictors were temperature, radiation, precipitation, soil moisture, humidity during the previous growing season between May and September and leaf senescence dates of the previous year, with random intercepts among species and sites. In addition, we quantified and compared the effects of climate variables of the previous growing season on leaf unfolding dates using boosted regression tree, an ensemble statistical learning method^67^, which has been widely applied in ecological modeling and prediction^68,69^. Because radiation and soil moisture data were only available since 1980, we selected phenology and climate datasets between 1984 and 2015 to perform the linear mixed-effects model and boosted regression tree. Linear mixed-effects model fitting was conducted using the *lme4* package^70^of R^64^. Significance testing of the fixed effects terms was done using the Satterthwaite method incorporated into the *lmerTest* package^71^ of R^64^, where *P* values less than 0.05 were considered significant. We performed the boosted regression trees using the *gbm* package^72^ in R^64^, where 10-fold cross validation was used to determine the optimal number of iterations.

### Effect of past climate change on spring phenology

Following Fu et al.^17^, we assessed the effects of past climate warming on spring phenology. First, we calculated the mean T_GS_ across all the 2,300 sites in Europe from 1951 to 2015. Using a 15-year smoothing window, we identified the coldest and warmest periods: 1955-1969 and 2000-2014 over the past 60 years. We calculated the S_T_ of leaf unfolding in response to the T_GS_ during the two periods for each species at each site. One-way analysis of variance (ANOVA) was used to test the difference in the S_T_ of leaf unfolding during 1955-1969 and 2000-2014.

### Structural equation modeling

We used a structural equation model (SEM) to analyze the relationships between climate, GPP_max_ and SOS from the 39 flux sites. The climate variables in the structural equation model included temperature, radiation, soil moisture, CO_2_ and precipitation during previous growing season. Because the daily GPP started to increase from DOY 120, peaking at DOY 180, then decreased until DOY 300 (Fig. S2), the period between May and September was also selected as the growing season. This is also consistently with the period of growing season identified by the dates of leaf unfolding and leaf senescence in PEP725 network. The SEM was fitted using the *lavaan* package^73^ in R^64^.

All data analyses were conducted using R version 4.0.3^64^.

## Results

Temperature sensitivity (S_T_, change in days per degree Celsius), is often used to describe the response of plant phenology to warmer temperatures. We calculated the S_T_ of spring phenology based on T_GS_ and dates of spring leaf unfolding obtained from PEP725 network, and start of season (SOS) metrics extracted from PhenoCam, and GIMMS NDVI_3g_ images (see Methods). The calculated S_T_ of spring phenology based on three datasets is shown in Fig. 2. Using the PEP725 network, the mean S_T_ of leaf unfolding across nine temperate tree species between 1951 and 2015 was −2.50 days·°C^-1^ (Fig. 2a). This suggested that a warmer previous growing season advanced leaf unfolding dates. The S_T_ was negative across all selected nine temperate tree species (Fig. 2b). The response of *Quercus robur* to T_GS_ were the strongest, with an average of −2.82 days·°C^-1^, significantly stronger than those of *Tilia cordata* (−1.04 days·°C^-1^) and *Tilia platyphyllos* (−1.16 days·°C^-1^).

**Fig. 2.**
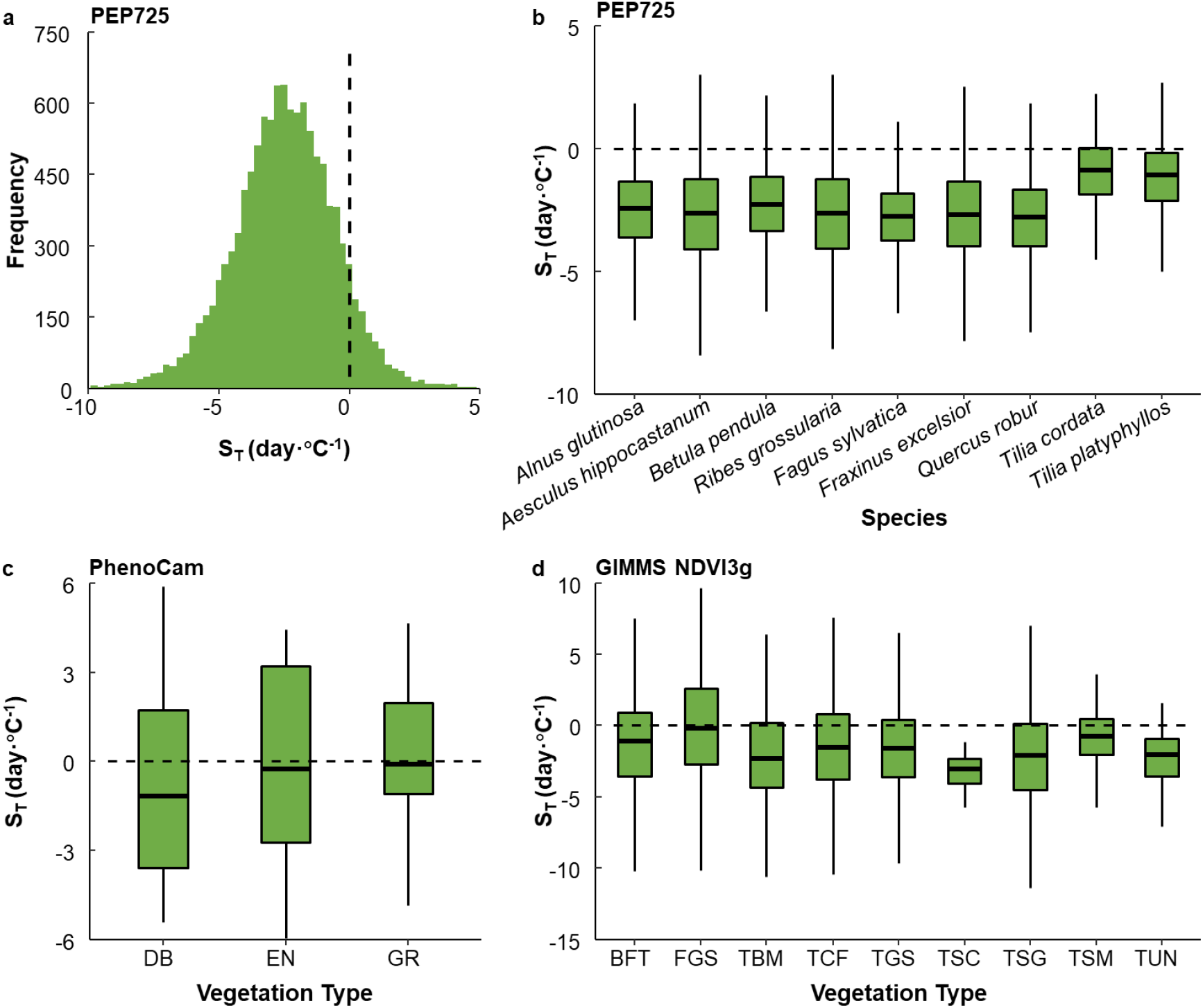
Temperature sensitivities (S_T_, change in days per degree Celsius) of spring phenology in response to increasing temperature during previous growing season. The calculated S_T_ was based on (**a, b**) records of spring leaf unfolding for nine temperate tree species at 2,300 sites in Europe, and phenological metrics extracted from (**c**) the PhenoCam network and (**d**) the GIMMS NDVI_3g_ products for different biomes. DB, EN and GR in (**c**) represents deciduous broad-leaved forests, evergreen forests and grasslands, respectively. In (**d**), the biomes included Boreal Forests/Taiga (BFT), Flooded Grasslands & Savannas (FGS), Temperate Broadleaf & Mixed Forests (TBM), Temperate Conifer Forests (TCF), Temperate Grasslands, Savannas & Shrublands (TGS), Tropical & Subtropical Coniferous Forests (TSC), Tropical &Subtropical Grasslands, Savannas & Shrublands (TSG), Tropical & Subtropical Moist Broadleaf Forest (TSM) and Tundra (TUN). The black dash lines indicate when the S_T_ is equal to zero.

In addition to temperature, spring phenology has been reported to be influenced by other climate variables and autumn phenology. We used a linear mixed effects model to exclude these co-variate effects and further examined the effects of T_GS_ on spring leaf unfolding. We consistently observed that leaf unfolding dates were advanced by increasing temperature by an average of −2.67 days·°C^-1^ (Table S2). Using boosted regression tree, we found the temperature had the strongest effect on leaf unfolding dates (84.67%), followed by radiation (6.33%), soil moisture (3.93%), precipitation (3.32%), humidity (1.75%) (Fig. S1).

Our PEP725 results were corroborated by PhenoCam and remote sensing data. Specifically, we observed a negative effect of T_GS_ on SOS in deciduous broad-leaved forests, evergreen forests and grasslands using phenological metrics extracted from the PhenoCam network between 2000 and 2018 (Fig. 2c). According to the calculated S_T_, the SOS in response to warming of the previous growing season was the strongest in deciduous broad-leaved forests, followed by evergreen forests and grasslands (Fig. 2c). Using the phenology metrics extracted from remote sensing dataset between 1982 and 2014, we also observed that increasing T_GS_ advanced SOS across different vegetation types in the Norther Hemisphere (Fig. 2d). Among all vegetation types, the S_T_ of the Tundra was the lowest, followed by Temperate Broadleaf & Mixed Forests and Savannas & Shrublands (Fig. 2d).

To test whether earlier spring phenology was driven by increased photosynthetic carbon assimilation, we further examined the relationship between daily maximum photosynthetic rate (GPP_max_) of the previous growing season and SOS between 1992 and 2014 using FLUXNET data. We found that the timing of SOS showed a significant negative correlation with the GPP_max_ during the growing season between 1992 and 2014 (correlation coefficient = −0.36, *P*<0.01, Fig. 3a). This suggested that spring phenology tended to occur earlier with the increased photosynthetic carbon assimilation during previous growing season. To further test the carbon-driven hypothesis, we constructed a structural equation model (SEM) that included climate variables, GPP_max_ and SOS (Fig. 3b). We found that spring phenology (SOS) was advanced by increased GPP_max_ (slope = −2.331, *P*<0.001). In addition, the effect of temperature on GPP_max_ was the strongest (slope = 0.319, *P*<0.001), followed by soil moisture (slope = 0.167, *P*<0.001), while radiation (slope = 0.005, *P*>0.05), CO_2_ (slope = 0.002, *P*>0.05) and precipitation (slope = 0.001, *P*>0.05) almost had no effects on GPP_max_. The detailed statistics of the SEM are listed in Table S3.

**Fig. 3.**
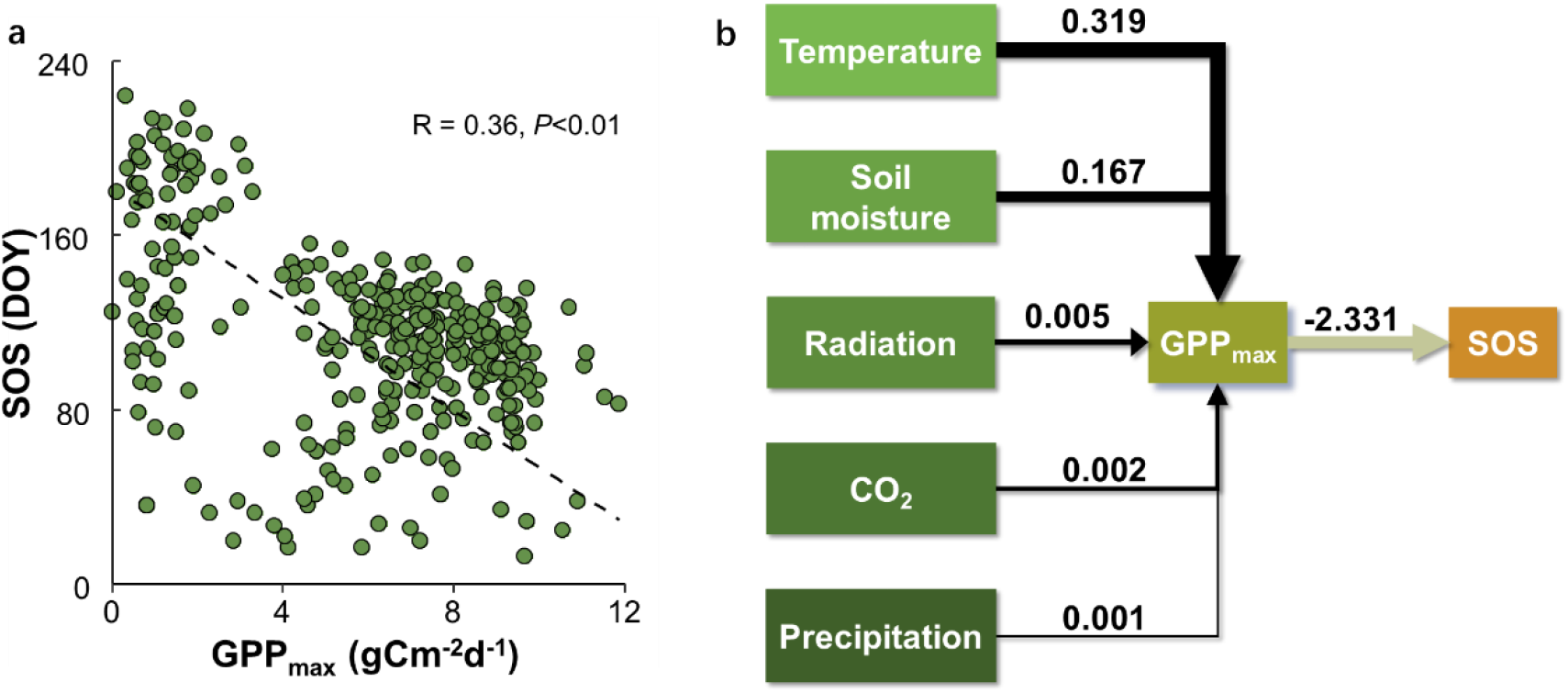
(**a**) Relationship between spring phenology (SOS) and GPP_max_ and (**b**) the constructed structural equation model using the data of 39 FLUXNET sites between 1992 and 2014. The black dash line represents the fitted linear regression line (SOS = 182.38−12.88×GPP_max_). The used variables in the structural equation model included climate variables (temperature, radiation, soil moisture, CO_2_ and precipitation), SOS and GPP_max_.

To examine the potential effects of climate warming on leaf unfolding, we used the PEP725 dataset to calculate and compare the S_T_ between the coldest and the warmest 15-year periods: 1955-1969 and 2000-2014, respectively (Figs. 4 and S3). We found that the S_T_ of leaf unfolding decreased by 63.1% from −1.76 ± 0. 04 days·°C^-1^ during 1955-1969 to −0.65 ± 0. 04 during 2000-2014 (Fig. 4a). Between 1955 and 1969, the S_T_ of early-successional species is −2.37 days·°C^-1^ and −1.23 days·°C^-1^ for late successional species. Between 2000 and 2014, S_T_ of the early-and late-successional species were −0.13 days·°C^-1^ and –0.92 days·°C^-1^, respectively. The S_T_ of the early successional species decreased more from the coldest to the warmest periods (−2.24 ± 0.15 days·°C^-1^) than that of late successional species (−0.31 ± 0.16 days·°C^-1^) (Figs. 4b and S3).

**Fig. 4.**
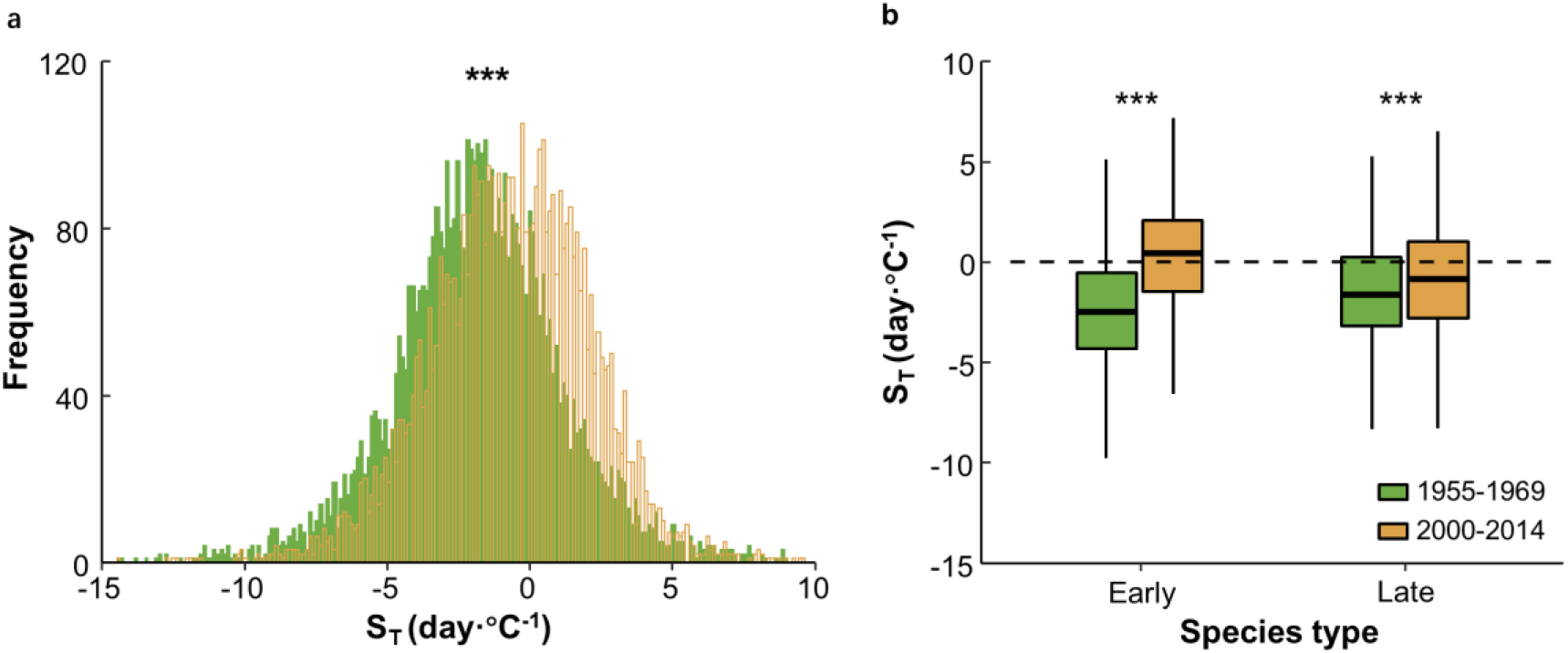
(**a**) Distributions of temperature sensitivities (S_T_, change in days per degree Celsius) of leaf unfolding during the coldest (1955-1969) and the warmest (2000-2014) periods and (**b**) differences of S_T_ between early- and late-successional species during these two periods. The calculated S_T_ was based on the temperature during previous growing season and leaf unfolding dates obtained from the PEP725 database. The length of each box indicates the interquartile range, the horizontal line inside each box the median, and the bottom and top of the box the first and third quartiles respectively. The asterisks indicate a significant difference in the S_T_ 1955-1969 and 2000-2014 (*P*<0.001). The black dashed horizontal line indicates when the S_T_ is equal to zero.

## Discussion

Global warming advances budbreak and leafing worldwide^21,74–77^. Using three long-term and large-scale phenological datasets, we show that warmer temperatures of the previous growing season drive earlier phenology in the following spring in the Northern Hemisphere. We also find that warming increased photosynthetic carbon assimilation, suggesting a physiological mechanism by which global warming is triggering earlier spring phenology (Fig. 5).

**Fig. 5.**
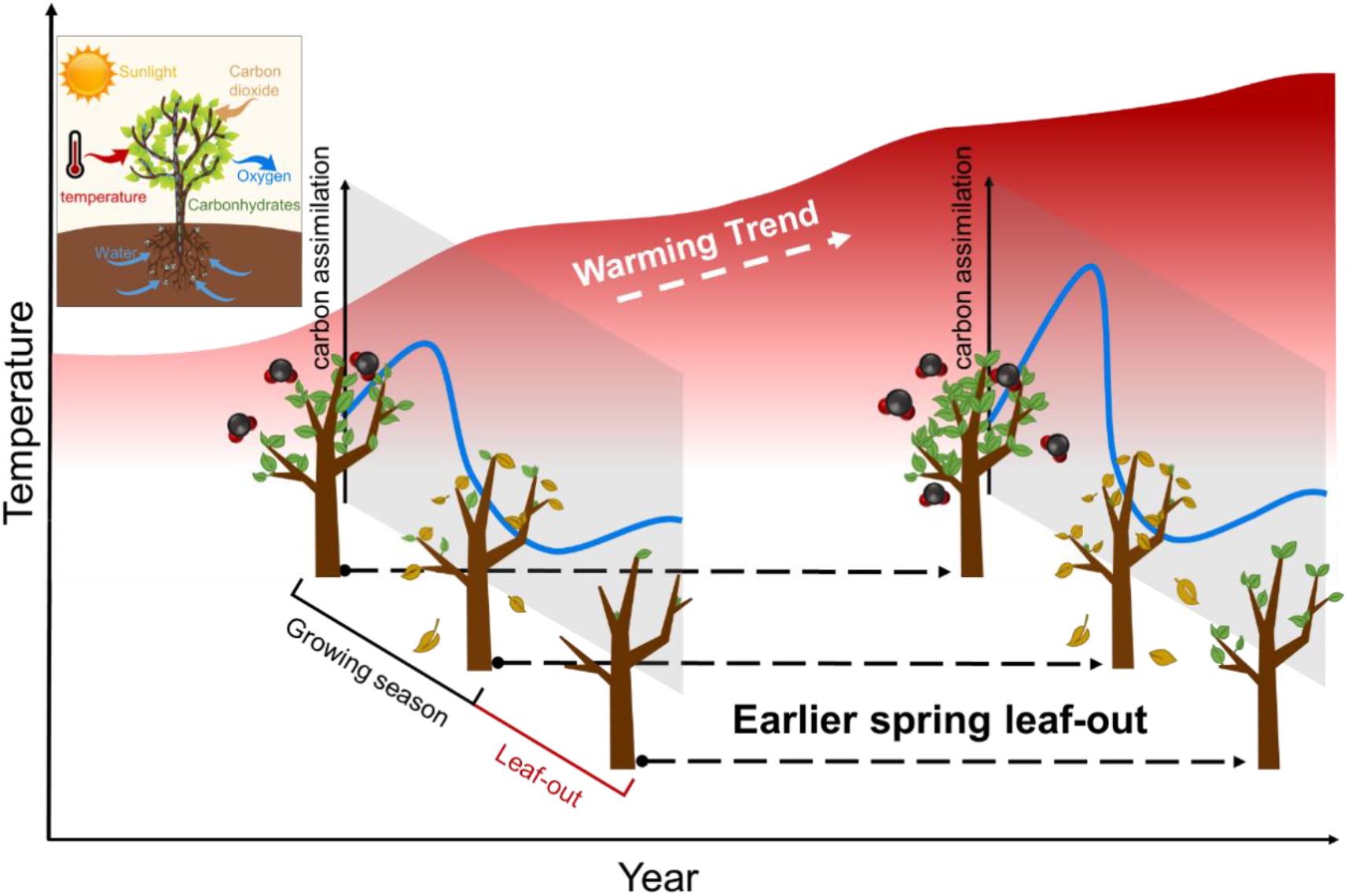
A schematic diagram of the earlier leaf-out in response to warming during previous growing season. Warmer temperatures during the previous growing season drivers earlier spring leaf-out by increasing photosynthetic carbon assimilation.

In deciduous tree species, carbon gained through photosynthesis is often stored in the form of non-structural carbohydrates (NSC-soluble carbohydrates and starch), which supports the growth of buds and leaves in the following spring before newly grown leaves can supply photosynthesis^78–80^. For instance, 95% of starches stored in the branches of *Fagus sylvatica* and *Quercus petraea* were consumed when spring bud-break occurred^79^. Needle growth of *Larix gmelinii* in spring drew nearly 50% of the carbohydrates fixed in the previous year^81,82^. Phloem girdling showed that deficient carbon storage can significantly delay the timing of spring budbreak and reduce bud size^27^.

During winter dormancy, temperate tree species also need to store sufficient carbohydrates prior to leaf senescence for respiration to maintain baseline functions and protect cells from frost damage and ensure survival^83,84^. Therefore, warmer temperatures in the previous growing season may advance spring phenology by increasing carbon storage, supported by the negative correlations between spring phenology and maximum photosynthetic rate in the previous year.

Recently, Zani et al.^32^ has reported that increased carbon assimilation during the growing season drives earlier autumn leaf senescence in temperate ecosystems. When leaf senescence occurred earlier, trees advanced the endodormancy^5,85^. In this context, the requirement of chilling units may be also fulfilled earlier. As a result, earlier autumn phenology facilitates an earlier spring phenology^86^. Therefore, increased carbon assimilation may directly drive autumn phenology, and, in turn, influence spring phenology. In our analyses, we excluded the co-variate effect of autumn phenology and isolated the effect of temperature of the previous growing season on leaf unfolding. The relationship was negative, confirming our hypothesis that increased carbon assimilation of previous season triggers an earlier spring phenology.

We observed that early-successional species showed a stronger response to the warming during growing season compared to late-successional species. In addition to temperature, spring phenology is also under photoperiodic control^87^. Because photoperiod remains stable regardless of climate change, plants are expected to show relatively conservative climatic responses when they rely on photoperiod to determine spring phenology. However, photoperiod sensitivities often vary among species^87^. For example, late-successional species are reported to have a higher photoperiod sensitivity compared to early-successional species^87,88^. The higher photoperiod sensitivity of late-successional species may, therefore, explain their conservative climatic responses compared to early-successional species ^87,89^.

Recent studies have reported that the warming-induced earlier spring phenology has slowed down over the past decades^21,90,91^. Fu et al.^17^ reported that S_T_ of leaf unfolding decreased by 40% from 4.0 ± 1.8 days·°C^-1^ during 1980-1994 to 2.3 ± 1.6 days·°C^-1^ during 1999-2013. The observed declining effect of warming on spring phenology is generally considered a result of chilling reduction in winter^92^. However, the carbon-driven earlier spring phenology is also slowing down in recent decades, especially for early-successional species as found here. Duffy et al.^12^ showed that the mean temperatures in the warmest quarter passed the optimal for photosynthesis over the past decade, with a sharp declining photosynthesis. The increased heat and water stress of the last decades may lead to a spreading growth decline of forests^93–95^. Therefore, the observed decline in the S_T_ may involve reductions in carbon assimilation by heat waves and/or drought events under global warming^96,97^.

## Conclusion

Despite the warming-induced spring phenology observed worldwide, the underlying causes and physiological mechanisms still remain unclear. In this study, we used multiple long-term and large-scale datasets to provide evidence that spring phenology is advanced by warmer temperatures of the previous growing season. Correspondingly, we observed that leaf unfolding occurred earlier under enhanced maximum photosynthetic capability. These findings suggest that an increased carbon assimilation under global warming could be involved in the observed earlier leafing of trees. In addition, we observed a decline in the carry-over effect of growing-season warming on spring phenology resulted likely from the reduced photosynthetic carbon assimilation by heat and water stress under global warming. With an increase in projected drought frequency under warming scenarios^93,98^, we expect that temperate trees will slow down the advancement of spring phenology. This may reduce the strength of forest carbon sinks under future climate conditions^17^. Our study provides new insights into the warming-induced change in spring phenology under global climate change to predict spring phenology and vegetation-atmosphere feedbacks under future climatic scenarios.

## Supporting Information

**Table S1.** List of the 9 temperate species selected from the PEP725 phenological network.

| Number | Latin name | Successional type |
| --- | --- | --- |
| 1 | *Aesculus hippocastanum* L. | Early |
| 2 | *Betula pendula* Roth | Early |
| 3 | *Alnus glutinosa* (L.) Gaertn. | Early |
| 4 | *Ribes grossularia* L. | Early |
| 5 | *Fraxinus excelsior* L. | Late |
| 7 | *Fagus sylvatica* L. | Late |
| 7 | *Quercus robur* L. | Late |
| 8 | *Tilia cordata* Mill. | Late |
| 9 | *Tilia platyphyllos* Scop. | Late |

**Table S2.**
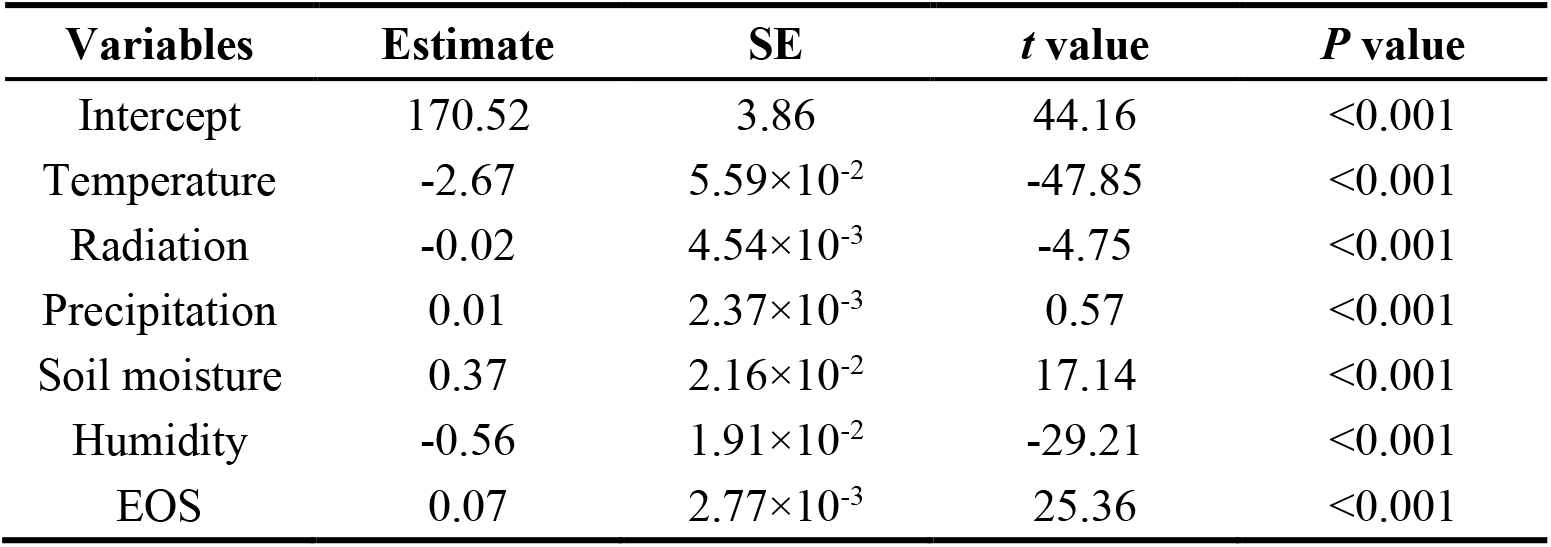
Results of linear mixed model that the effect of temperature during previous growing season on spring phenology (SOS) after excluding the influence of other climatic factors (radiation, precipitation, soil moisture, humidity) and autumn phenology (EOS).

**Table S3.** Statistics of the structural equation models (SEMs). To display model performance, we calculated the Comparative Fit Index (CFI) and the root-mean square error (RMSEA).

| **Statistics of the structural equation models (SEMs)** |  |  |  |  |  |  |
| --- | --- | --- | --- | --- | --- | --- |
| **Left-hand side** | **Option** | **Right-hand side** | **Estimate** | **SE** | **Z value** | **P value** |
| GPP<sub>max</sub> | ~ | Temperature | 0.319 | 0.033 | 9.569 | <0.001 |
| GPP<sub>max</sub> | ~ | Soil moisture | 0.167 | 0.026 | 6.347 | <0.001 |
| GPP<sub>max</sub> | ~ | Radiation | 0.005 | 0.004 | 1.179 | >0.05 |
| GPP<sub>max</sub> | ~ | CO<sub>2</sub> | 0.002 | 0.007 | 0.325 | >0.05 |
| GPP<sub>max</sub> | ~ | Precipitation | 0.001 | 0.152 | 0.004 | >0.05 |
| SOS | ~ | GPP<sub>max</sub> | -2.331 | 0.551 | -4.229 | <0.001 |
| CFI = 0.89; RMSEA = 0.20 |  |  |  |  |  |  |

**Fig. S1.**
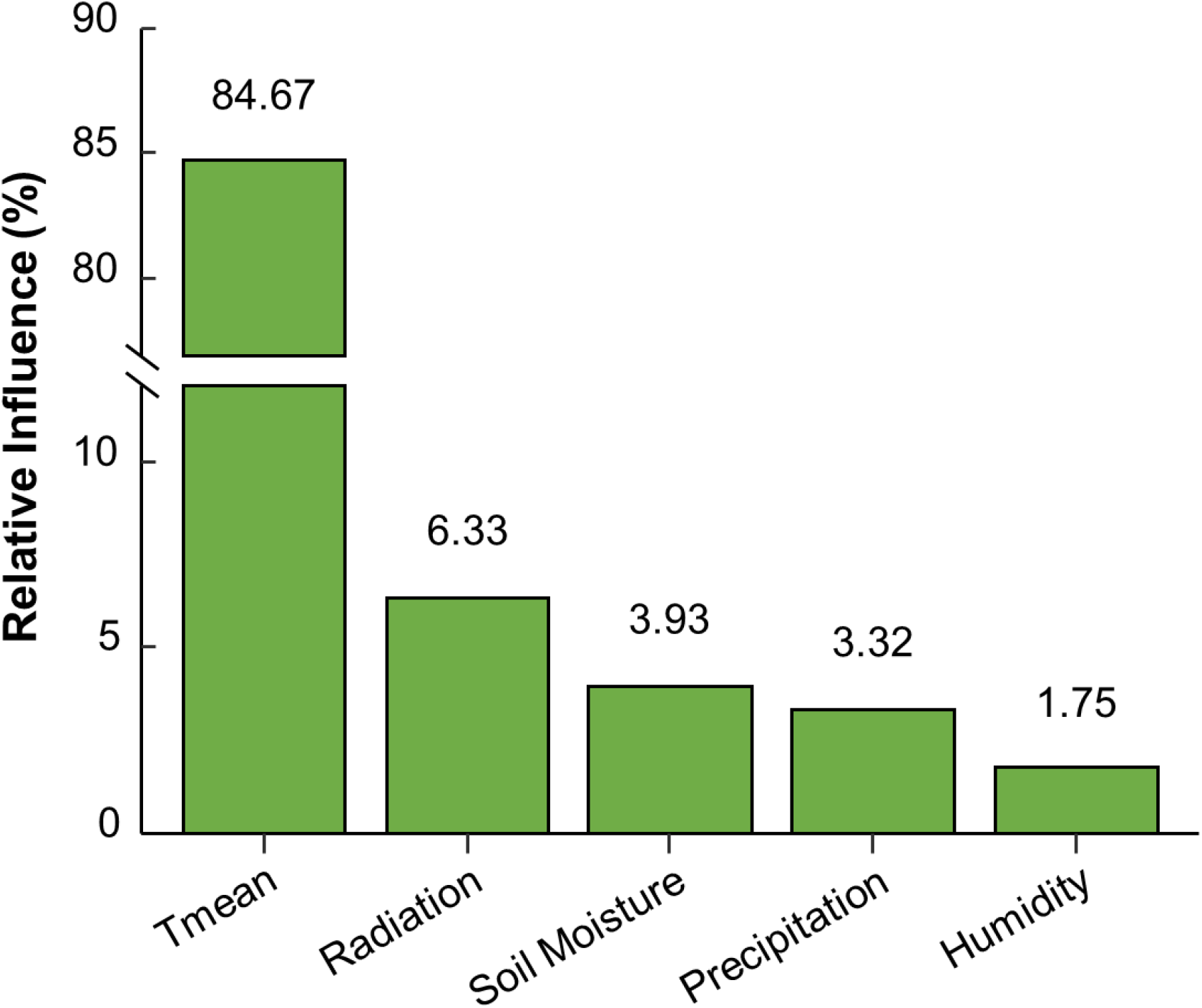
The relative influence of climatic factors during previous growing season on spring leaf unfolding between 1984 and 2015 obtained from the PEP725 database.

**Fig. S2.**
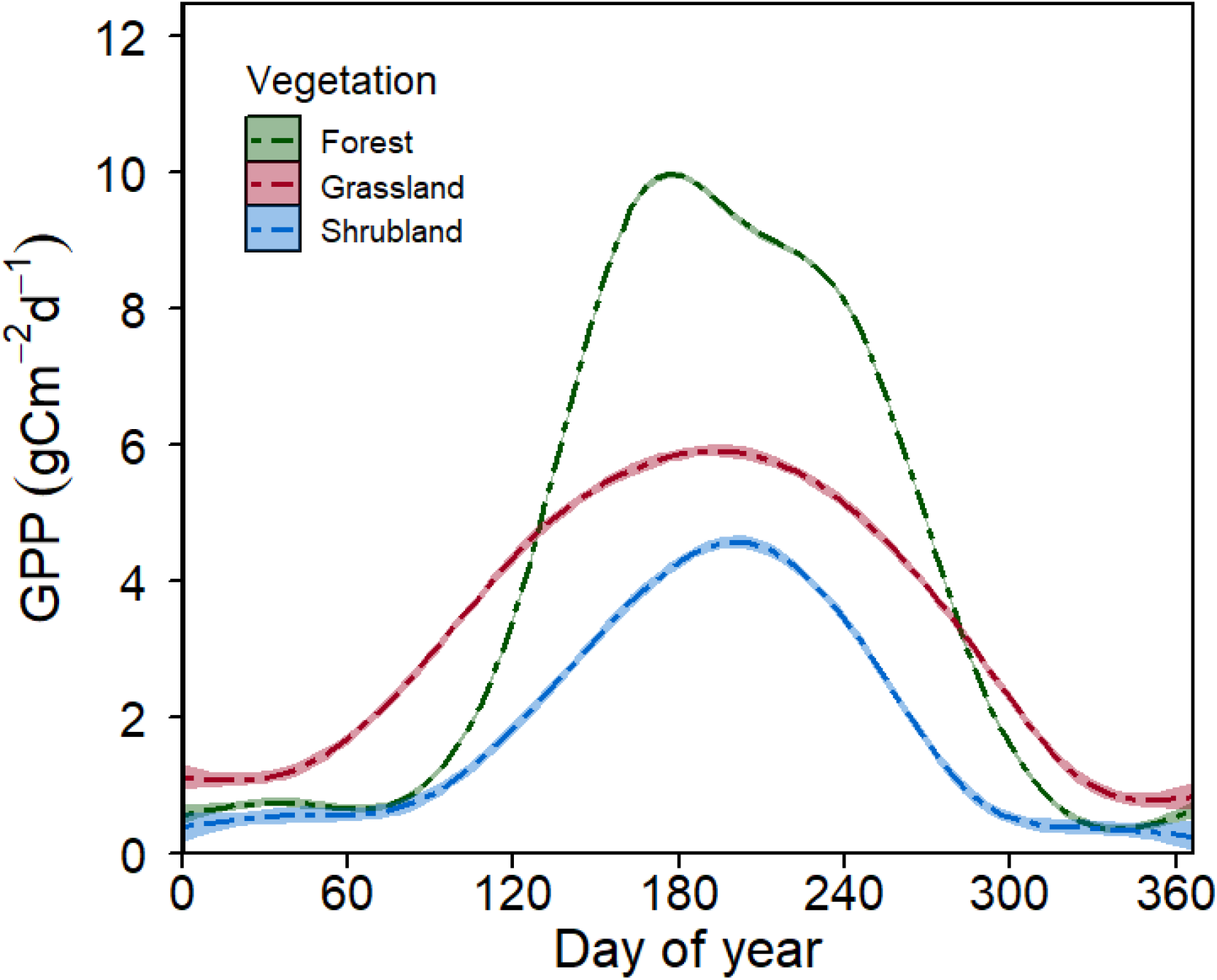
Daily gross primary productivity (GPP) changes in three vegetation types based on FLUXNET.

**Fig. S3.**
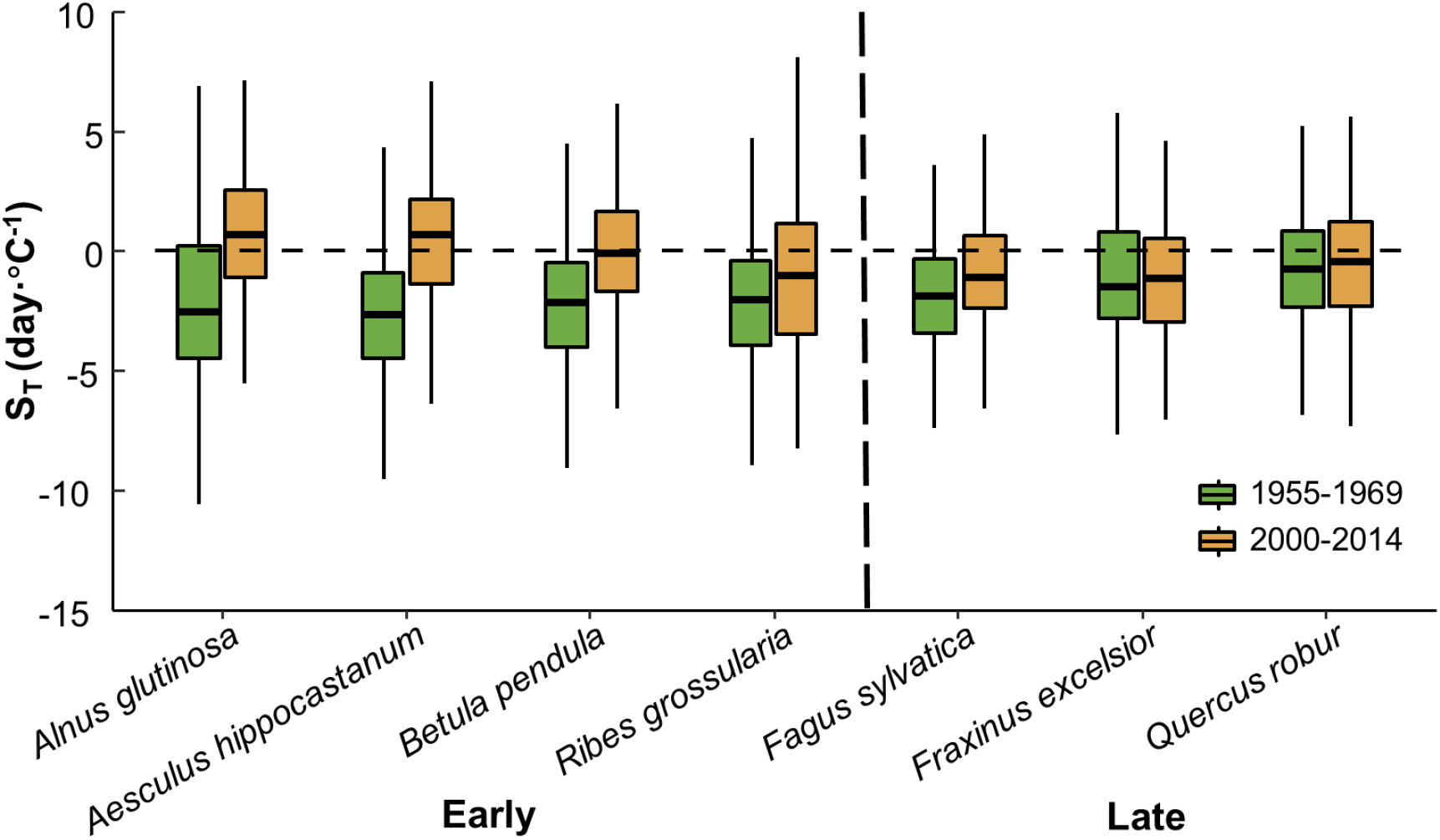
Temperature sensitivities (S_T_, change in days per degree Celsius) of leaf unfolding in early- and late-successional species during 1955-1969 and 2000-2014. The calculated S_T_ was based on the temperature during previous growing season and leaf unfolding dates obtained from the PEP725 database. The length of each box indicates the interquartile range, the horizontal line inside each box the median, and the bottom and top of the box the first and third quartiles respectively. The black dashed horizontal line indicates when the S_T_ is equal to zero.

## Notes

### Competing Interest Statement

The authors have declared no competing interest.

### Summary of Updates

The version of the manuscript modified the grammar and legend.

## References

1. Gao, M. et al.. Divergent changes in the elevational gradient of vegetation activities over the last 30 years. Nat. Commun. 10, 2970 (2019).

2. Koen Hufkens et al. Ecological impacts of a widespread frost event following early spring leaf-out. Glob. Chang. Biol. 18, 2365–2377 (2012).

3. Kari Saikkonen et al. Climate change-driven species’ range shifts filtered by photoperiodism. Nat. Clim. Chang. 2, 239–242 (2012).

4. Zeng, H., Jia, G., & Howard Epstein. Recent changes in phenology over the northern high latitudes detected from multi-satellite data. Environ. Res. Lett. 6, 45508–45518 (2011).

5. Marc Estiarte & Josep Peñuelas. Alteration of the phenology of leaf senescence and fall in winter deciduous species by climate change: effects on nutrient proficiency. Glob. Chang. Biol. 21, 1005–1017 (2015).

6. Richardson, A. et al.. Climate change, phenology, and phenological control of vegetation feedbacks to the climate system. Agric. For. Meteorol. 169, 156–173 (2013).

7. Josep Penuelas, This Rutishauser, & Iolanda Filella. Phenology Feedbacks on Climate Change. Science 324, 887–888 (2009).

8. Fu, Y. H. et al.. Three times greater weight of daytime than of night-time temperature on leaf unfolding phenology in temperate trees. New Phytol. 212, 590–597 (2016).

9. Annette Menzel et al. European phenological response to climate change matches the warming pattern. Glob. Chang. Biol. 12, 1969–1976 (2006).

10. Peuelas, J. & Iolanda Filella. Responses to a Warming World. Science 294, 793–795 (2001).

11. Piao, S. et al.. Leaf onset in the northern hemisphere triggered by daytime temperature. Nat. Commun. 6, 6911 (2015).

12. Duffy, K. A. et al.. How close are we to the temperature tipping point of the terrestrial biosphereã Sci. Adv. 7, eaay1052 (2021).

13. Keenan, T. F., Richardson, A. D. & Hufkens, K. On quantifying the apparent temperature sensitivity of plant phenology. New Phytol. 225, 1033–1040 (2020).

14. Niu, S., Fu, Y., Gu, L. & Luo, Y. Temperature Sensitivity of Canopy Photosynthesis Phenology in Northern Ecosystems. Phenology: An Integrative Environmental Science 503–519 doi:10.1007/978-94-007-6925-0_27 (2013).

15. Meineke, E. K., Davis, C. C. & Davies, T. J. Phenological sensitivity to temperature mediates herbivory. Glob. Chang. Biol. 27, 2315–2327 (2021).

16. Huanjiong Wang, Quansheng Ge, This Rutishauser, Yuxiao Dai, & Junhu Dai. Parameterization of temperature sensitivity of spring phenology and its application in explaining diverse phenological responses to temperature change. Sci. Rep. 5, 8833 (2015).

17. Fu, Y. H. et al.. Declining global warming effects on the phenology of spring leaf unfolding. Nature 526, 104–107 (2015).

18. Carol K Augspurger. Reconstructing patterns of temperature, phenology, and frost damage over 124 years: Spring damage risk is increasing. Ecology 94, 41–50 (2013).

19. Jochner, S., Beck, I., Behrendt, H., Traidl-Hoffmann, C. & Menzel, A. Effects of extreme spring temperatures on phenology: a case study from Munich and Ingolstadt. Clim. Res. 12, 101–112 (2010).

20. Meng, F., Zhang, L. irong, Zhang, Z., Jiang, L. & Wang, Y. Enhanced spring temperature sensitivity of carbon emission links to earlier phenology. Sci. Total Environ. 745, 140999 (2020).

21. Güsewell, R., Furrer, R., Gehrig, R. & Pietragalla, B. Changes in temperature sensitivity of spring phenology with recent climate warming in Switzerland are related to shifts of the preseason. Glob. Chang. Biol. 23, 5189–5202 (2017).

22. Wang, T. et al.. The influence of local spring temperature variance on temperature sensitivity of spring phenology. Glob. Chang. Biol. 20, 1473–1480 (2014).

23. Huang, J.-G. et al.. Intra-annual wood formation of subtropical Chinese red pine shows better growth in dry season than wet season. Tree Physiol. 38, 1225–1236 (2018).

24. Knowles, J. F., Scott, R. L., Biederman, J. A., Blanken, P. D. & Barron-Gafford, G. A. Montane forest productivity across a semi-arid climatic gradient. Glob. Chang. Biol. 26, 6945–6958 (2020).

25. Strimbeck, G. R., Trygve, D. K., Paul, G. S. & Paula, F. M. Dynamics of low-temperature acclimation in temperate and boreal conifer foliage in a mild winter climate. Tree Physiol. 28, 1365–1374 (2008).

26. Smart, D. R. Exposure to elevated carbon dioxide concentration in the dark lowers the respiration quotient of Vitis cane wood. Tree Physiol. 24, 115–120 (2004).

27. Roxas, A. A., Orozco, J., Guzmán-Delgado, P., & Zwieniecki, M. A. Spring phenology is affected by fall non-structural carbohydrates concentration and winter sugar redistribution in three Mediterranean nut tree species. Tree Physiol. 1-14 (2021).

28. Palacio, S., Maestro, M., & G Montserrat-Martí. Seasonal dynamics of non-structural carbohydrates in two species of mediterranean sub-shrubs with different leaf phenology. Environ. Exp. Bot. 59, 34–42 (2007).

29. Fierravanti, A., Rossi, S., Kneeshaw, D., De Grandpré, L. & Deslauriers, A. Low Non-structural Carbon Accumulation in Spring Reduces Growth and Increases Mortality in Conifers Defoliated by Spruce Budworm. Front. For. Glob. Chang. 2, (2019).

30. Fajstavr, M., Giagli, K., Vavrcik, H., Gryc, V. & Urban, J. The effect of stem girdling on xylem and phloem formation in Scots pine. Silva Fennica 51, 1760 (2017).

31. Fajstavr, M. et al.. The cambial response of Scots pine trees to girdling and water stress. Iawa Journal 41, 159–185 (2020).

32. Zani, D., Crowther, T. W., Mo, L., Renner, S. S. & Zohner, C. M. Increased growing-season productivity drives earlier autumn leaf senescence in temperate trees. Science 370, 1066–1071 (2020).

33. Lin, Y.-S., Medlyn, B. E. & Ellsworth, D. S. Temperature responses of leaf net photosynthesis: the role of component processes. Tree Physiol. 32, 219–231 (2012).

34. Smith, N. G. & Dukes, J. S. Plant respiration and photosynthesis in global-scale models: incorporating acclimation to temperature and CO2. Glob. Chang. Biol. 19, 45–63 (2013).

35. Dusenge, M. E., Duarte, A. G. & Way, D. A. Plant carbon metabolism and climate change: elevated CO2 and temperature impacts on photosynthesis, photorespiration and respiration. New Phytol. 221, 32–49 (2019).

36. Huang, M. et al.. Air temperature optima of vegetation productivity across global biomes. Nat. Ecol. Evol. 3, 772–779 (2019).

37. Smith, N. G., Lombardozzi, D., Tawfik, A., Bonan, G. & Dukes, J. S. Biophysical consequences of photosynthetic temperature acclimation for climate. J. Adv. Model. 9, 536–547 (2017).

38. Saxe, H., Cannell, M. G. R., Johnsen, Ø., Ryan, M. G. & Vourlitis, G. Tree and forest functioning in response to global warming. New Phytol. 149, 369–399 (2001).

39. Bigras, F. J. & Bertrand, A. Responses of Picea mariana to elevated CO2 concentration during growth, cold hardening and dehardening: phenology, cold tolerance, photosynthesis and growth. Tree Physiol. 26, 875–888 (2006).

40. M.Ewa, J. & Reinhart, C. Effects of elevated atmospheric CO2 on phenology, growth and crown structure of Scots pine (Pinus sylvestris) seedlings after two years of exposure in the field. Tree Physiol. 19, 289–300 (1999).

41. Penuelas & J. Phenology. Responses to a warming world. Science 294, 793–795 (2001).

42. Rebecca, S.-D. et al.. Divergent carbon cycle response of forest and grass-dominated northern temperate ecosystems to record winter warming. Glob. Chang. Biol. 26, 1519–1531 (2019).

43. Templ, B. et al.. Pan European Phenological database (PEP725): a single point of access for European data. Int. J. Biometeorol. 62, 1109–1113 (2018).

44. Leys, C., Ley, C., Klein, O., Bernard, P. & Licata, L. Detecting outliers: Do not use standard deviation around the mean, use absolute deviation around the median. J. Exp. Soc. Psychol. 49, 764–766 (2013).

45. Brown, M. T., Campbell, D. E., De Vilbiss, C. & Ulgiati, S. The geobiosphere emergy baseline: A synthesis. Ecol. Modell. 339, 92–95 (2016).

46. Richardson, A. D. et al.. Tracking vegetation phenology across diverse North American biomes using PhenoCam imagery. Sci. Data. 5, 180028 (2018).

47. Klosterman, S. T. et al.. Evaluating remote sensing of deciduous forest phenology at multiple spatial scales using PhenoCam imagery. Biogeosciences 11, 4305–4320 (2014).

48. Zhang, Y. et al.. Seasonal and interannual changes in vegetation activity of tropical forests in Southeast Asia. Agric. For. Meteorol. 224, 1–10 (2016).

49. Pinzon, J. E. & Tucker, C. J. A Non-Stationary 1981–2012 AVHRR NDVI3g Time Series. Remote Sens. 6, 6929–6960 (2014).

50. Wang, X. et al.. No trends in spring and autumn phenology during the global warming hiatus. Nat. Commun. 10, 2389 (2019).

51. Wang, X. et al.. Validation of MODIS-GPP product at 10 flux sites in northern China. Int. J. Remote Sens. 34, 587–599 (2013).

52. Julien, Y. & Sobrino, J. A. Global land surface phenology trends from GIMMS database. Int. J. Remote Sens. 30, 3495–3513 (2009).

53. Zhang, X. et al.. Monitoring vegetation phenology using MODIS. Remote Sens. Environ. 84, 471–475 (2003).

54. Pastorello, G. et al.. The FLUXNET2015 dataset and the ONEFlux processing pipeline for eddy covariance data. Sci. Data. 7, 225 (2020).

55. Kalman & Dan. A singularly valuable decomposition: The SVD of a matrix. The College Mathematics Journal 39, 2233–2241 (1996).

56. Huang, K. et al.. Enhanced peak growth of global vegetation and its key mechanisms. Nat. Ecol. Evol. 2, 1897–1905 (2018).

57. Tang, Y., Xu, X., Zhou, Z., Qu, Y. & Sun, Y. Estimating global maximum gross primary productivity of vegetation based on the combination of MODIS greenness and temperature data. Ecol. Inform. 63, 101307 (2021).

58. Xia, J. et al.. Joint control of terrestrial gross primary productivity by plant phenology and physiology. Proc. Natl. Acad. Sci. U.S.A. 112, 2788–2793 (2015).

59. Hu, Z. et al.. Joint structural and physiological control on the interannual variation in productivity in a temperate grassland: A data-model comparison. Glob. Chang. Biol. 24, 2965–2979 (2018).

60. Richardson, A. D. et al.. Influence of spring and autumn phenological transitions on forest ecosystem productivity. Philos. Trans. R. Soc. Lond. B. Biol. Sci. 365, 3227–3246 (2010).

61. Wu, C. et al.. Interannual variability of net carbon exchange is related to the lag between the end-dates of net carbon uptake and photosynthesis: Evidence from long records at two contrasting forest stands. Agric. For. Meteorol. 164, 29–38 (2012).

62. Cornes, R. C., Schrier, G. van der, Besselaar, E. J. M. van den & Jones, P. D. An Ensemble Version of the E-OBS Temperature and Precipitation Data Sets. J. Geophys. Res. Atmos. 123, 9391–9409 (2018).

63. Hijmans, R. J. raster: Geographic Data Analysis and Modeling. (2020).

64. R Core Team. R: A Language and Environment for Statistical Computing. (2020).

65. Suonan, J., Classen, A. T., Sanders, N. J. & He, J.-S. Plant phenological sensitivity to climate change on the Tibetan Plateau and relative to other areas of the world. Ecosphere 10, e02543 (2019).

66. Wang, C., Cao, R., Chen, J., Rao, Y. & Tang, Y. Temperature sensitivity of spring vegetation phenology correlates to within-spring warming speed over the Northern Hemisphere. Ecol. Indic. 50, 62–68 (2015).

67. Elith, J., Leathwick, J. R. & Hastie, T. A working guide to boosted regression trees. J Anim. Ecol. 77, 802–813 (2008).

68. Davis, K. T. et al.. Wildfires and climate change push low-elevation forests across a critical climate threshold for tree regeneration. Proc. Natl. Acad. Sci. U.S.A. 116, 6193–6198 (2019).

69. Lemm, J. U. et al.. Multiple stressors determine river ecological status at the European scale: Towards an integrated understanding of river status deterioration. Glob. Chang. Biol. 27, 1962–1975 (2021).

70. Bates, D., Mächler, M., Bolker, B. & Walker, S. Fitting Linear Mixed-Effects Models Using lme4. J. Stat. Softw. 67, 1–48 (2015).

71. Kuznetsova, A., Brockhoff, P. B. & Christensen, R. H. B. lmerTest Package: Tests in Linear Mixed Effects Models. J. Stat. Softw. 82, 1–26 (2017).

72. Greenwell, B., Boehmke, B., Cunningham, J. & Developers, G. B. M. gbm: Generalized Boosted Regression Models. (2020).

73. Rosseel, Y. lavaan: An R Package for Structural Equation Modeling. 48 (2012).

74. Asch & Rebecca, G. Climate change and decadal shifts in the phenology of larval fishes in the California Current ecosystem. Proc. Natl. Acad. Sci. U.S.A. 112, 4065–4074 (2015).

75. RaphaSaarn. False estimates of the advance of spring. Nature 600 (2001).

76. Wheeler, H. C., Hye, T. T., Schmidt, N. M., Svenning, J. C. & Forchhammer, M. C. Phenological mismatch with abiotic conditions—implications for flowering in Arctic plants. Ecology 96, 775–787 (2015).

77. Yu, H., Luedeling, E. & Xu, J. Winter and spring warming result in delayed spring phenology on the Tibetan Plateau. Proc. Natl. Acad. Sci. U.S.A. 107, 22151–22156 (2010).

78. Hartmann, H. & Trumbore, S. Understanding the roles of nonstructural carbohydrates in forest trees – from what we can measure to what we want to know. New Phytol. 211, 386–403 (2016).

79. Tamir, K., Yann, V., & H Günter. Coordination between growth, phenology and carbon storage in three coexisting deciduous tree species in a temperate forest. Tree Physiol. 36, 847–855 (2016).

80. Tixier, A., Gambetta, G. A., Godfrey, J., Orozco, J. & Zwieniecki, M. A. Non-structural Carbohydrates in Dormant Woody Perennials; The Tale of Winter Survival and Spring Arrival. Front. For. Glob. Chang. 2, 18 (2019).

81. Kagawa, A. & Maximov, S. Seasonal course of translocation, storage and remobilization of 13C pulse-labeled photoassimilate in naturally growing Larix gmelinii saplings. New Phytol. 171, 793–804 (2010).

82. Siegwolf et al. Examining the response of needle carbohydrates from Siberian larch trees to climate using compound-specific C-13 and concentration analyses. Plant Cell Environ. 38, 2340–2352 (2015).

83. Erica, F., Eduardo, F., Helen, B. & Eike, L. A Conceptual Framework for Winter Dormancy in Deciduous Trees. Agronomy 10, 241 (2020).

84. Majken, P., Brandt, A. U. & Lillie, A. Winter warming delays dormancy release, advances budburst, alters carbohydrate metabolism and reduces yield in a temperate shrub. Aob Plants 7, plv024 (2015).

85. Malyshev, A. V. Warming Events Advance or Delay Spring Phenology by Affecting Bud Dormancy Depth in Trees. Front. Plant Sci. 11, 856 (2020).

86. Liu, Q. et al.. Modeling leaf senescence of deciduous tree species in Europe. Glob. Chang. Biol. 26, (2020).

87. Basler, D. & Körner, C. Photoperiod sensitivity of bud burst in 14 temperate forest tree species. Agric. For. Meteorol. 165, 73–81 (2012).

88. Basler, D. & Körner, C. Photoperiod and temperature responses of bud swelling and bud burst in four temperate forest tree species. Tree Physiol. 34, 377–388 (2014).

89. Körner, C. & Basler, D. Phenology Under Global Warming. Science 327, 1461 (2010).

90. Daphné, A. et al.. Warmer winters reduce the advance of tree spring phenology induced by warmer springs in the Alps. Agric. For. Meteorol. 252, 220–230 (2018).

91. Vitasse, Signarbieux, & YSH. Global warming leads to more uniform spring phenology across elevations. Proc. Natl. Acad. Sci. U.S.A. 155, (1004).

92. Clark, J. S., Salk, C., Melillo, J. & Mohan, J. Tree phenology responses to winter chilling, spring warming, at north and south range limits. Funct. Ecol. 28, 1344–1355 (2014).

93. Brzostek, E. R. et al.. Chronic water stress reduces tree growth and the carbon sink of deciduous hardwood forests. Glob. Chang. Biol. 20, 2531–2539 (2014).

94. Julio Camarero, J. et al. Forest Growth Responses to Drought at Short- and Long-Term Scales in Spain: Squeezing the Stress Memory from Tree Rings. Front. Ecol. Evol. 6, (2018).

95. Xu, P. et al.. Impacts of Water Stress on Forest Recovery and Its Interaction with Canopy Height. Int. J. Environ. Res. Public Health. 15, 1257 (2018).

96. Yashavanthakumar, K. J. et al.. Impact of heat and drought stress on phenological development and yield in bread wheat. Plant Physiol. Rep. 26, 357–367 (2021).

97. Choukri, H. et al.. Heat and Drought Stress Impact on Phenology, Grain Yield, and Nutritional Quality of Lentil (Lens culinaris Medikus). Front. Nutr. 7, (2020).

98. Zhou, S., Zhang, Y., Williams, A. P. & Gentine, P. Projected increases in intensity, frequency, and terrestrial carbon costs of compound drought and aridity events. Sci. Adv. 5, eaau5740 (2019).

